# Inactivation of a non-canonical gp130 signaling arm attenuates chronic systemic inflammation and multimorbidity induced by a high-fat diet

**DOI:** 10.1101/2024.04.08.588362

**Authors:** Youngjoo Lee, Arijita Sarkar, Jade Tassey, Jonathan N. Levi, Siyoung Lee, Nancy Q. Liu, Andrew C. Drake, Jenny Magallanes, Una Stevic, Jinxiu Lu, Dawei Ge, Hanhan Tang, Tadiwanashe Mkaratigwa, Fangzhou Bian, Ruzanna Shkhyan, Michael Bonaguidi, Denis Evseenko

**Author notes:** Correspondence to: Denis Evseenko MD, PhD. Associate Professor of Orthopaedic Surgery, Stem Cell Research and Regenerative Medicine. USC, Los Angeles, CA, 90033, USA. 1450 Biggy St. NRT4509, Los Angeles, CA 90033.

## Abstract

Interleukin-6 (IL-6) is a major pro-inflammatory cytokine for which the levels in plasma demonstrate a robust correlation with age and body mass index (BMI) as part of the senescence-associated secretory phenotype. IL-6 cytokines also play a crucial role in metabolic homeostasis and regenerative processes, primarily via the canonical STAT3 pathway. Thus, selective modulation of IL-6 signaling may offer a unique opportunity for therapeutic interventions. Recently, we discovered that a non-canonical signaling pathway downstream of tyrosine (Y) 814 within the intracellular domain of gp130, the IL-6 co-receptor, is responsible for the recruitment and activation of SRC family of kinases (SFK). Mice with constitutive genetic inactivation of gp130 Y814 (F814 mice) show accelerated resolution of inflammatory response and superior regenerative outcomes in skin wound healing and posttraumatic models of osteoarthritis. The current study was designed to explore if selective genetic or pharmacological inhibition of the non-canonical gp130-Y814/SFK signaling reduces systemic chronic inflammation and multimorbidity in a high-fat diet (HFD)-induced model of accelerated aging. F814 mice showed significantly reduced inflammatory response to HFD in adipose and liver tissue, with significantly reduced levels of systemic inflammation compared to wild type mice. F814 mice were also protected from HFD-induced bone loss and cartilage degeneration. Pharmacological inhibition of gp130-Y814/SFK in mice on HFD mirrored the effects observed in F814 mice on HFD; furthermore, this pharmacological treatment also demonstrated a marked increase in physical activity levels and protective effects against inflammation-associated suppression of neurogenesis in the brain tissue compared to the control group. These findings suggest that selective inhibition of SFK signaling downstream of gp130 receptor represents a promising strategy to alleviate systemic chronic inflammation. Increased degenerative changes and tissue senescence are inevitable in obese and aged organisms, but we demonstrated that the systemic response and inflammation-associated multi-morbidity can be therapeutically mitigated.

## Introduction

The mechanisms involved in systemic chronic inflammatory syndrome, also known as “inflammaging” or “non-genetic accelerated aging syndrome”, are poorly understood and often associated with chronological aging, obesity, and chronic stress. This gap in knowledge limits the development of therapies used to manage inflammaging and associated multimorbidity that is marked by the accumulation of senescent cells combined with increased systemic release and persistence of pro-inflammatory factors, such as Interleukin 6 (IL-6) (1–3). Inflammaging in older adults is usually marked in the clinic by stable elevated levels of C-reactive protein (CRP) and monocyte chemoattractant protein-1 (MCP-1) (4,5) with no other symptoms of a specific inflammatory disorder such as an autoimmune condition or chronic infection. Elevation of CRP (especially above 3 mg/L) and increased body mass index (BMI) (above 30 kg/m^2^), are associated with increased risk of multi-organ failure and mortality in multiple studies conducted in different ages, ethnical and racial groups (6–9). Increased levels of MCP-1 in aging adults have been associated with decreases in memory, cardiovascular disease, and sarcopenia (4,10). IL-6 is one of the key factors regulating the senescence-associated secreted phenotype produced by senescent cells in degenerating tissues and in immune cells activated by this progressive multiorgan degeneration and atrophy(11–13). IL-6 is recognized as the key upstream activator of CRP production by the liver cells (14,15) and of MCP-1 production in multiple tissues (16,17)

All IL-6 family of cytokines signal through the transmembrane receptor gp130, which mediates downstream signaling via its intracellular residues that impact distinct biological processes. In our recent study, we have identified a signaling tyrosine 814 (Y814) residue within the gp130 intracellular domain that acts as a “stress sensor” responsible for triggering degenerative, and pathological changes in tissues via activation of SRC family of kinases (SFK) (18). Endogenous products arising from tissue senescence, which is associated with aging, may also cause maladaptive hyperactivation of this sensor, which is highly detrimental. CRISPR/Cas9 mice with a constitutively inactivated gp130^Y814^ (mutation of tyrosine (Y) 814 to phenylalanine (F) – “F814” mice exhibit substantially accelerated regenerative responses (18). Both genetic and drug-mediated inactivation of SFK recruitment to gp130 receptor result in enhanced resolution of inflammation and accelerated regenerative outcomes in several injury models (18).

However, the role of this novel regulatory mechanism in chronic systemic inflammation is not yet defined. Multiple studies have shown that an experimental high-fat diet (HFD) leads to chronic, low-grade systemic inflammation and progressive dysfunction of multiple organs and tissues. HFD causes osteoarthritis (OA) in mice (19,20), sarcopenia, osteoporosis, and multiple organ dysfunction, including endocrine organs, liver, kidney, and cardiac (reviewed in (21)). HFD also leads to a significant elevation of IL-6, MCP-1, and CRP levels in the systemic circulation (21–23). We hypothesized that the non-canonical gp130-Y814/SFK signaling was a novel mechanism implicated in the pathogenesis of chronic systemic inflammation and associated multimorbidity. We also predicted that selective targeting of this fundamental mechanism can improve the healthspan of animals with systemic chronic inflammation induced by HFD. The current study demonstrates that both genetic and drug-mediated disruption of SFK recruitment to gp130 results in the attenuation of systemic inflammation and improved functional capacity of multiple organs including the liver, musculoskeletal joints and bone, normalization of cholesterol metabolism, and increased physical activity in an accelerated aging model of mice fed with HFD.

## Results

### Genetic mutation of gp130^Y814^ ameliorates systemic inflammation induced by HFD

To investigate the role of gp130^Y814^ activation in a systemic chronic inflammatory condition, 2-month-old F814 and wild type (WT) C57BL/6J mice were placed on HFD for 10 months (Figure 1A, B). As expected, HFD increased the weight and fat content nearly 2-fold and significantly decreased bone mineral density compared to the normal diet (ND) group (Figure S1A). The HFD group exhibited adipocyte hyperplasia and a crown-like pattern of macrophages surrounding necrotic adipocytes, indicating the inflammatory changes in adipose tissue (Figure S1B). The livers of HFD-fed mice had an excess accumulation of lipid droplets, increased fibrosis, and macrophage infiltration, all of which are characteristics of non-alcoholic fatty liver disease (Figure S1C).

**Figure 1.**
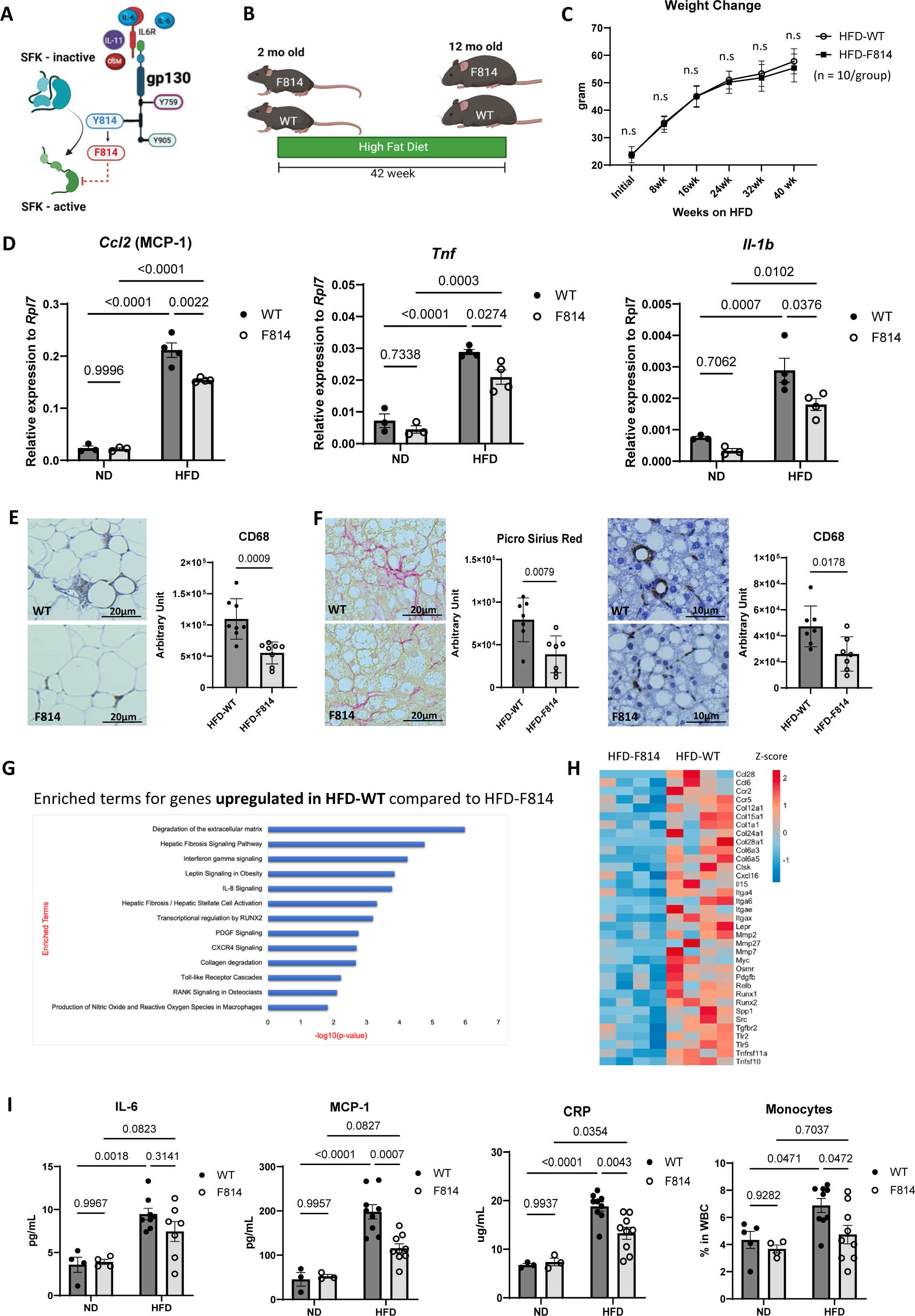
Gp130-Y814 mutant mice show reduced systemic inflammation induced by a high fat diet (HFD). (A) IL-6/gp130 activation of SRC family of kinases (SFKs) via gp130 Y814 residue. A point mutation of gp130 tyrosine (Y) 814 to phenylalanine (F) 814 inhibited SFK activation via IL-6/gp130 signaling. (B) 2-month-old F814 mice and wild type (WT) counterparts were administered HFD for 10 months. Mice were 12 months old by the end of the experiment. (C) Weight change of WT and F814 mice during HFD administration, n= 10 per group. Multiple unpaired t-test with FDR = 5% was used to compare each time point and q-values less than 0.05 were considered significant. (D) qPCR analysis of adipose tissue expression of *Ccl2* (MCP-1), *Tnf,* and *Il1b* relative to a housekeeping gene *Rpl7* in HFD-fed WT and F814 compared to normal diet (ND) counterparts. (E) Representative images of CD68 staining of adipose tissue of WT and F814 mice on HFD and quantitative analysis. (F) Representative images of Picro Sirius Red staining and CD68 staining of liver tissue of WT and F814 mice on HFD and quantitative analysis. (G) Enriched terms for genes upregulated in the liver of WT on HFD compared to F814 counterparts analyzed based on RNA-seq data. (H) Heat map showing differentially regulated genes (HFD-F814 vs HFD-WT) associated with inflammation and fibrosis in the liver from bulk RNA-seq. (I) IL-6, MCP-1, and CRP levels in serum measured by ELISA and monocyte ratio in peripheral blood analyzed from white blood cell (WBC) differential. Statistical analysis for (D), (H): 2-way ANOVA with multiple comparisons with Tukey, p-values less than 0.05 were considered significant; (E), (F): Two-tailed Student’s t-test, p-values less than 0.05 were considered significant.

F814 mutant mice on HFD gained a similar amount of weight and fat mass compared to WT (Figure 1C, S2A). We expected HFD to induce inflammatory responses in major metabolic organs, so we first evaluated the adipose tissue and the liver of F814 and WT mice. Although transcription of genes representative of adipose tissue-driven inflammation, *Ccl2* (MCP-1), *Tnf*, and *Il1b* was upregulated in both F814 and WT mice on a HFD compared to their ND counterparts, the mutant mice exhibited reduced upregulation compared to the WT (Figure 1D). Adipose tissue from F814 mice on HFD showed reduced macrophage infiltration (Figure 1E). Flow cytometry analysis of immune cells in adipose tissue from HFD mice revealed significantly fewer pro-inflammatory CD11b^+^/Ly6C^hi^ monocytes and CD11b^+^/F4/80^+^ macrophages in F814 than in WT (Figure S2B-C).

The livers of HFD-fed mice had an excessive accumulation of lipid droplets in both genotypes, but F814 mice showed reduced fibrosis and macrophage infiltration (Figure 1F). Consistent with this, bulk RNA sequencing (RNA-seq) analysis of livers showed an increase in inflammatory and fibrotic signaling in WT mice on HFD compared to F814 mice on HFD (Figure 1G). In particular, livers from WT on HFD had upregulated *Ccr2, Ccr5, Osmr, Spp1, Src, Tlr2,* and *Tlr5,* suggesting more macrophage infiltration and activated immune responses than in F814 (Figure 1H). Upregulation of *Col1a1, Ctsk, Itga4, Mmp2,* and *Pdgfb*, indicating fibrotic activity, was also attenuated in F814 on HFD than in WT (Figure 1H).

Since both the liver and adipose tissue are major contributors of secreted inflammatory mediators in obesity, we next tested if the levels of systemic inflammation differ between WT and F814 mice on HFD. Serum IL-6 levels were comparable between WT and mutant mice on HFD; however, F814 mice exhibited significantly lower serum levels of MCP-1, a classical monocyte recruiter that is required for the migration of monocytes from bone marrow, which is especially critical for chronic inflammatory conditions (24,25) (Figure 1I). The mutant had significantly lower levels of circulating monocytes, corroborating the lower MCP-1 level (Figure 1I). Neutrophil and lymphocyte ratio was not significantly different among the groups (Figure S2D). F814 mice on HFD also showed attenuated levels of CRP (Figure 1I). These results suggest that gp130^Y814^ residue may play a critical role in amplifying the pro-inflammatory signaling induced by HFD, which in turn promotes pathological changes and pro-inflammatory activity of the adipose tissue and liver, thereby increasing systemic inflammation levels.

### F814 mice demonstrate resistance to bone and cartilage loss induced by HFD

Previous studies have shown that chronic inflammation is one of the major drivers of bone and cartilage catabolism in various diseases (26–28). The DEXA scans did not show significant differences in bone mineral density between WT and F814 on HFD (Figure S2A), however, HFD-induced bone loss is reported to be in trabecular bones rather than cortical bones, which cannot be evaluated by the resolution of DEXA scans (29). Micro-computed tomography (µCT) was utilized to analyze bone mineral density of trabecular bone beneath the growth plate of proximal tibia (Figure 2A). While both WT and F814 mice on HFD lost trabecular bone density compared to the ND counterparts, F814 mice on HFD showed significant retention of trabecular bone compared to WT on HFD (Figure 2B).

**Figure 2.**
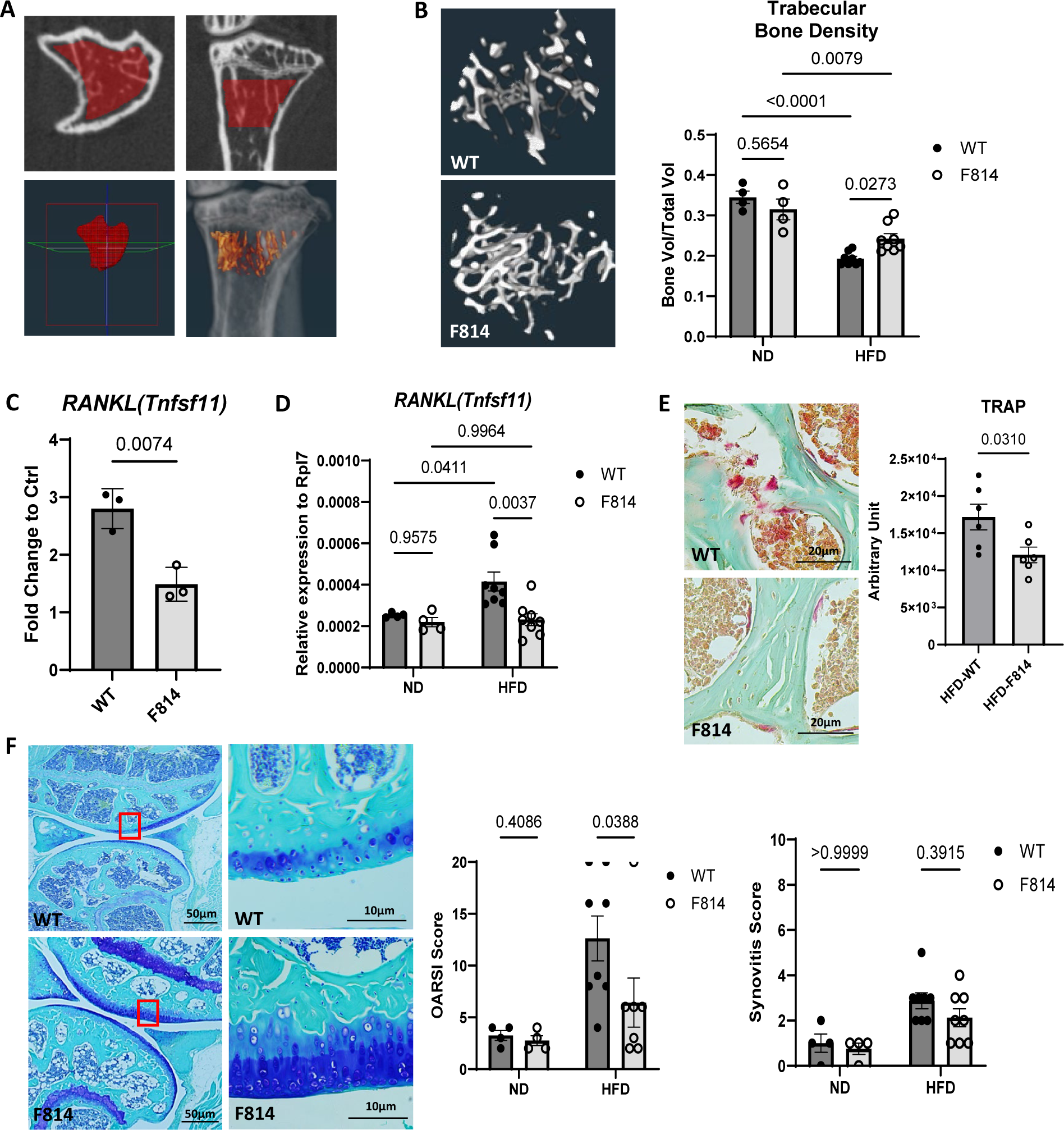
Gp130-Y814 mutant mice showed resistance to bone density loss and osteoarthritis induced by HFD. (A) Area of interest for analyzing trabecular bone density. Micro-CT scans of the trabecular bone beneath the growth plate of the proximal tibia were reconstructed in 3D. (B) Representative 3D constructed images of trabecular bone of WT and F814 mice on HFD and quantitative analysis on bone density. (C) Fold change of RANKL *(Tnfsf11)* expression levels of WT and F814 mesenchymal stromal cells (MSCs) stimulated with hyper-IL-6 (10ng/mL) compared to non-stimulated WT and F814 MSC controls. (D) Expression of RANKL *(Tnfsf11)* relative to a housekeeping gene *Rpl7* in bone marrow of HFD-fed WT and F814 compared to ND counterparts. (E) Representative images of TRAP staining (red) in trabecular bone (green) of WT and F814 mice on HFD and quantitative analysis. (F) Representative images of Toluidine Blue staining on joint articular cartilage from WT and F814 mouse knee and evaluation on OA and synovitis. OARSI score 0-4: No OA; 5-9: Mild OA; 10-14: Moderate OA; 15 and over: Severe OA. Synovitis score 1-4: low grade; 5-9: high grade. Multiple Mann-Whitney test with FDR = 10% was used for statistical analysis, q-values less than 0.05 were considered significant. Statistical analysis for (B), (D): 2-way ANOVA with multiple comparisons with Tukey correction, p-values less than 0.05 were considered significant; (C), (E): Two-tailed Student’s t-test, p-values less than 0.05 were considered significant.

To determine whether the higher bone density was a result of enhanced osteogenesis or reduced bone resorption under chronic inflammation, we evaluated the osteogenic and osteoclastogenic potential of F814 derived cells *in vitro*. Mesenchymal stromal cells (MSCs) derived from bone marrow were differentiated into osteocytes and bone matrix production was quantified by Alizarin Red staining. F814 cells produced comparable amounts of bone matrix as WT counterparts regardless of the inflammatory conditions induced by hyper-IL-6 or Oncostatin M (OSM), a pro-inflammatory IL-6 family cytokine (Figure S2E). F814 monocytes also differentiated into osteoclasts at similar levels as WT cells when treated with the same concentration of Receptor activator of nuclear factor-κB ligand (RANKL) (data not shown). RANKL is secreted by osteoblasts, osteocytes, bone marrow stromal cells, and immune cells when stimulated by pro-inflammatory cytokines including IL-6 (30,31). F814 MSCs expressed reduced levels of RANKL*(Tnfsf11)* gene upon hyper-IL-6 stimulation compared to WT counterparts (Figure 2C). In agreement with this, non-cultured total bone marrow cells from F814 mice on HFD expressed significantly lower levels of RANKL (*Tnfsf11)* compared to WT on HFD (Figure 2D). TRAP staining also showed significantly reduced number of osteoclasts in F814 on HFD compared to WT counterparts (Figure 2E). Collectively, the disruption of gp130-Y814 signaling serves as a protective factor against bone density loss induced by a high-fat diet through a reduction in RANKL production, ultimately ameliorating osteoclast activity and bone resorption induced by systemic chronic inflammation.

Long term administration of HFD is reported to induce early onset or significantly worsened progression of OA in mice (32). Morphometric analysis of the joints showed that F814 mice’ articular cartilage was largely protected from OA induced by HFD compared to WT mice. The majority of mutant mice on HFD had mild or minimal symptoms of OA, while WT counterparts developed mild to severe OA (Figure 2F). SFK activation is reported to be critical for cartilage degeneration in OA, and our group has demonstrated that F814 cartilage explants were resistant to OSM-induced degeneration (18,33). Mild synovitis, a characteristic of OA induced by low-grade chronic inflammation, was present in the majority of mice in both WT and F814 groups, although F814 mice exhibited a lower trend in synovitis score (Figure 2F).

### Pharmacological inhibition of gp130/SFK signaling reduces inflammatory responses in mice on HFD

Our previous studies showed that pharmacological inhibition of SFK signaling downstream of gp130 leads to anti-inflammatory and anti-fibrotic effects in acute local injuries in small and large animal models (18). Previously reported SFK recruitment inhibitor R805, developed by our group, had poor water solubility and was only suitable for local applications (18). R159, an analog of R805, was developed with an improved ADME profile to enhance water solubility for better systemic administration (Figure S3A). Similar to R805, R159 inhibited IL-6 cytokine-stimulated recruitment and activation of SFK by gp130 (Figure 3A, Figure S3B-C). To evaluate the effects of gp130^Y814^/SFK inactivation on systemic chronic inflammation, 1-year-old C57BL/6 mice were placed on HFD and administered R159 or vehicle (Veh) twice a week via intraperitoneal injection for 12 weeks (Figure 3B). For this pharmacological intervention study, older mice were used to closely model progression of obesity-induced multimorbidity in middle-aged humans. We also conducted additional experiments to evaluate its effects of the drug therapy to enhance our understanding of its therapeutic potential in age-associated chronic conditions.

**Figure 3.**
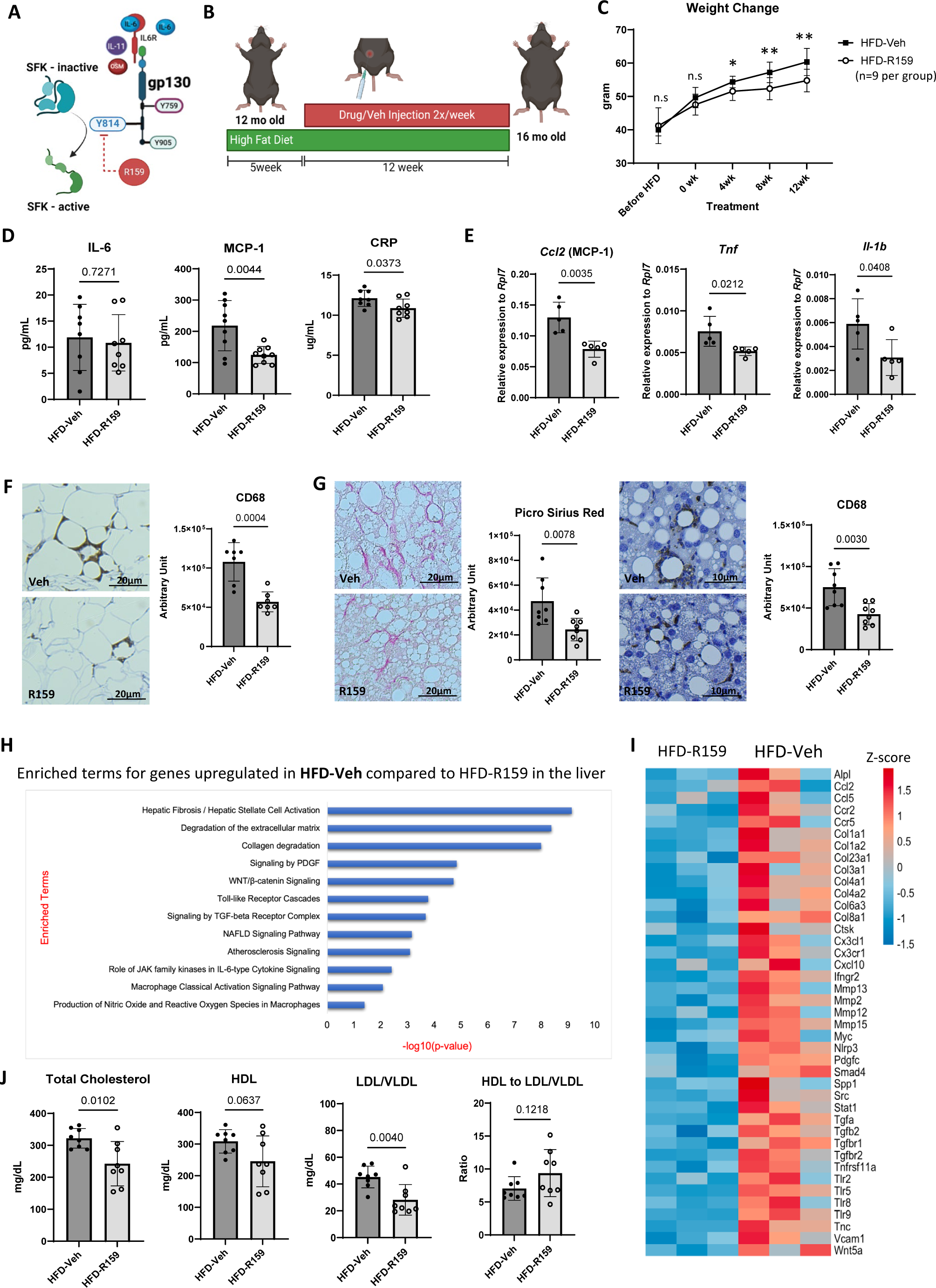
Gp130 modulating small molecule R159 reduced systemic chronic inflammation induced by aging and HFD. (A) IL-6/gp130 activation of Src Family Kinase (SFK) via Y814 residue was inhibited by a small molecule drug R159. (B) Experimental scheme to assess the effect of R159 on HFD-driven systemic chronic inflammation. 12-month-old mice were administered HFD for 5 weeks to induce systemic inflammation before the drug or vehicle (control) treatment began. Drug or vehicle was injected twice a week intraperitoneally for 12 weeks while the mice were continued on HFD. Mice were 16 months old by the end of the experiment. (C) Weight change of drug or vehicle-treated mice during HFD administration, n= 9 per group. Multiple unpaired t-tests with FDR = 5% were used to compare each time point, q-values less than 0.05 were considered significant. (D) Systemic levels of IL-6, MCP-1, and CRP in HFD mice treated either with vehicle or R159 measured by ELISA. (E) qPCR analysis of adipose tissue expression of *Ccl2* (MCP-1), *Tnf,* and *Il1b* relative to a housekeeping gene *Rpl7* in vehicle or R159 treated mice on HFD. (F) Representative images of CD68 staining of adipose tissue of vehicle or R159 treated mice on HFD and quantitative analysis. (G) Representative images of Picro Sirius Red staining and CD68 staining of liver tissue of vehicle or R159 treated mice on HFD and quantitative analysis. (H) Enriched terms for genes upregulated in the liver of vehicle group compared to R159 group analyzed based on RNA-seq data. (I) Heat map showing differentially regulated genes (HFD-R159 vs HFD-Veh) associated with inflammation and fibrosis in the liver from bulk RNA-seq. (J) Total cholesterol, High Density Lipoprotein (HDL), Low Density Lipoprotein (LDL), and Very Low Density Lipoprotein (VLDL) in the serum of HFD mice treated with vehicle or R159 were measured using a Cholesterol Assay kit. Statistical analysis for (D), (E), (F), (G), (J): Two-tailed Student’s t-test, p-values less than 0.05 were considered significant.

Notably, mice treated with R159 gained less weight compared to the Veh group (Figure 3C). R159-treatment mirrored the F814 mutant group’s systemic inflammatory marker patterns, significantly lowering systemic levels of MCP-1 and CRP without altering IL-6 levels compared to the Veh-treated group (Figure 3D). The drug had no discernible adverse effects during the long-term treatment and organ histology revealed no anomalies (data not shown).

Similar to genetic inactivation of gp130^Y814^ signaling, R159 treatment attenuated expression of *Ccl2* (MCP-1), *Tnf,* and *Il1b* in adipose tissue induced by HFD (Figure 3E). Adipose tissue from R159 treated mice on HFD showed reduced macrophage infiltration compared to Veh treated mice (Figure 3F). R159 treated mice liver tissue also exhibited reduced fibrosis and macrophage infiltration compared to the Veh counterparts (Figure 3G). RNA-seq analysis of the liver showed an increase in inflammatory and fibrotic pathways in mice treated with Veh on HFD compared to the R159 group (Figure 3H). Similar to the liver RNA-seq results of F814 vs WT on HFD, Veh treated mice had upregulated *Ccr2, Ccr5, Cx3cr1, Spp1, Src, Tlr2,* and *Tlr5,* suggesting increased macrophage presence and immune response, and upregulated *Col1a1, Ctsk, Mmp2,* and *Pdgfc*, indicating more fibrosis activity compared to R159 counterparts (Figure 3I). The RNA-seq results also suggested enrichment of atherosclerosis-associated genes in the Veh group compared to the R159 group (Figure 3H). Serum lipid analysis revealed that R159 treatment significantly reduced total cholesterol and LDL/VLDL, resulting in higher HDL to LDL/VLDL ratio (Figure 3J).

Taken together, R159 treatment effectively mitigated systemic inflammatory responses of major metabolic organs and weight gain in mice on HFD and provided substantial protection to liver function. R159 treatment did not exhibit any adverse effects or histological anomalies, demonstrating its potential as a therapeutic intervention with minimal side effects.

### R159 mitigates degenerative changes induced by HFD in musculoskeletal tissues

The attenuation of systemic chronic inflammation by R159 treatment in HFD-fed mice revealed a multifaceted therapeutic potential, significantly reducing adverse effects of HFD. Bone loss induced by HFD was even more severe in older mice but was significantly mitigated by R159 treatment (Figure 4A). The drug treatment group showed reduced osteoclasts marked by TRAP staining in trabecular bone sections and lower expression of the RANKL (*Tnfsf11)* gene in the bone marrow compared to the Veh counterparts (Figure 4B-C). Additionally, while the vast majority of vehicle-treated mice on HFD exhibited more advanced OA, R159-treated HFD mice showed trends of milder osteoarthritis and lower synovitis score, although the effects were not statistically significant (Figure 4D).

**Figure 4.**
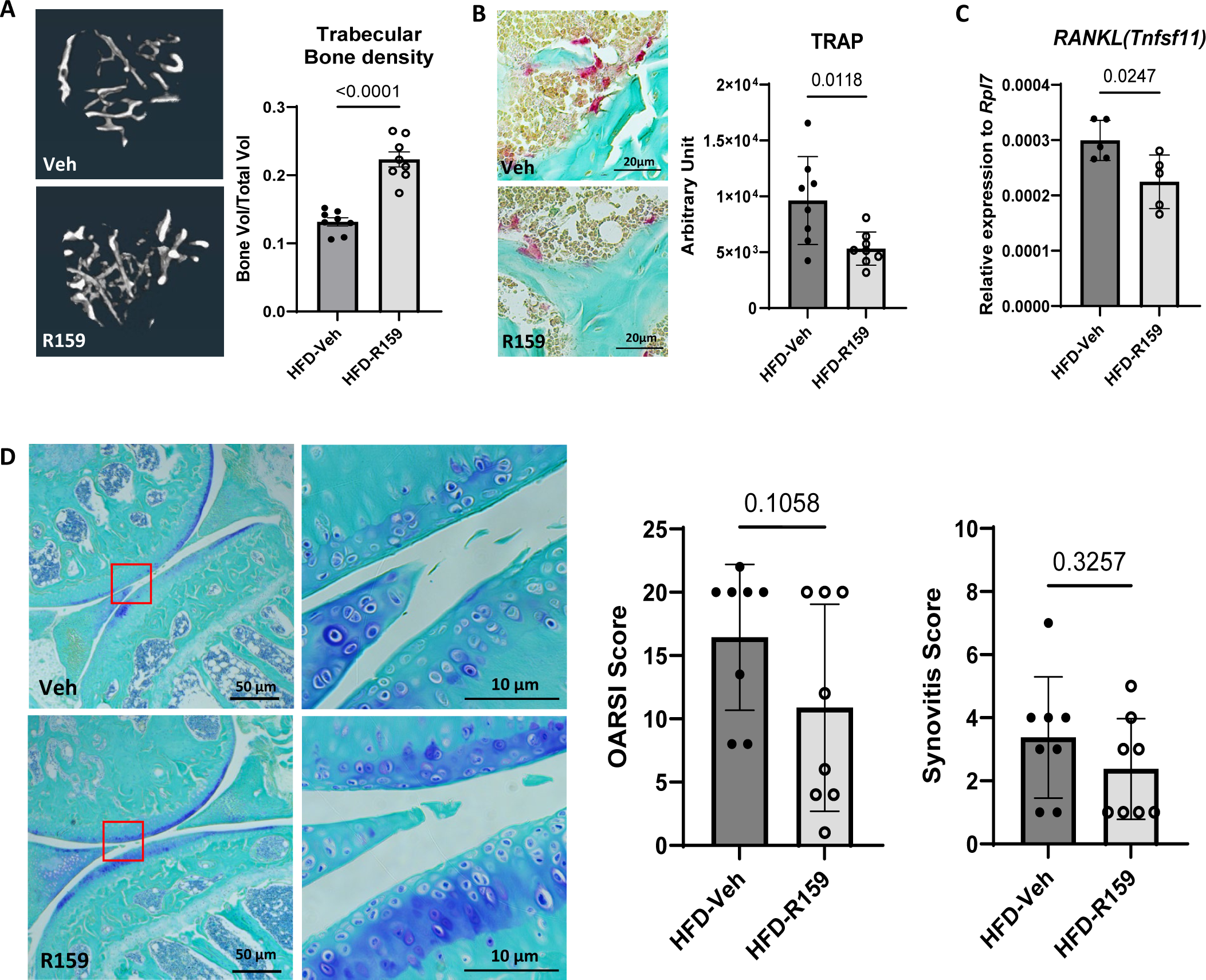
R159 treatment protected mice from musculoskeletal tissue degeneration induced by HFD. (A) Representative 3D constructed images of trabecular bone of vehicle or R159 treated mice on HFD and quantitative analysis on bone density. (B) Representative images of TRAP staining (red) in trabecular bone (green) of vehicle or R159 treated mice on HFD and quantitative analysis. (C) Expression of RANKL *(Tnfsf11)* relative to a housekeeping gene *Rpl7* in bone marrow of vehicle or R159 treated mice on HFD. (D) Representative images of Toluidine Blue staining on joint articular cartilage staining and evaluation on osteoarthritis and synovitis. Two-tailed Mann-Whitney test was used for statistical analysis and p-values less than 0.05 were considered significant. Statistical analysis for (A), (B), (C): Two-tailed Student’s t-test, p-values less than 0.05 were considered significant.

### Inhibition of gp130/SFK signaling mitigates inflammation-induced decline in neurogenesis

Exhaustion of stem cells is known to be a major hallmark of aging leading to reduced regenerative potential in various tissue (34). We next tested if R159 has a protective effect in several stem cell niches in mice with HFD-induced accelerated aging. Although R159 treatment showed no effect on hematopoietic and skin stem cell pools (Figure S3D-E), R159 provided significant protective effects against the inflammation-induced reduction of neurogenesis. Neural stem and progenitor cells diminish with age and obesity (35), likely contributing to the cognitive decline observed in aged, obese animals (36). The hippocampus, a critical brain region for memory formation and learning, harbors neural stem cells that continuously replenish neurons into adulthood (37). At 12 months of age, we observed there was no significant difference in the neural stem cell (NSC) number and neurogenesis even when the mice were put on HFD (Figure S3F). However, WT mice on HFD showed substantial decrease in proliferating NSCs, suggesting inflammation-induced niche disturbance (Figure S3F). F814 mice on HFD, on the other hand, maintained NSC proliferation levels similar to ND controls (Figure S3F). In mice, the decrease in NSC number, neurogenesis, and NSC proliferation starts as early as 6 months of age, with a substantial decline occurring around 12 to 18 months (36). We too observed a significant decline in the NSC pool, neurogenesis, and NSC proliferation between 12-month-old and 16-month-old WT mice (Figure S3G). Importantly, 16-month-old R159-treated mice on HFD showed significantly enhanced NSC maintenance and substantially increased neurogenesis compared to Veh-treated counterparts (Figure 5A).

**Figure 5.**
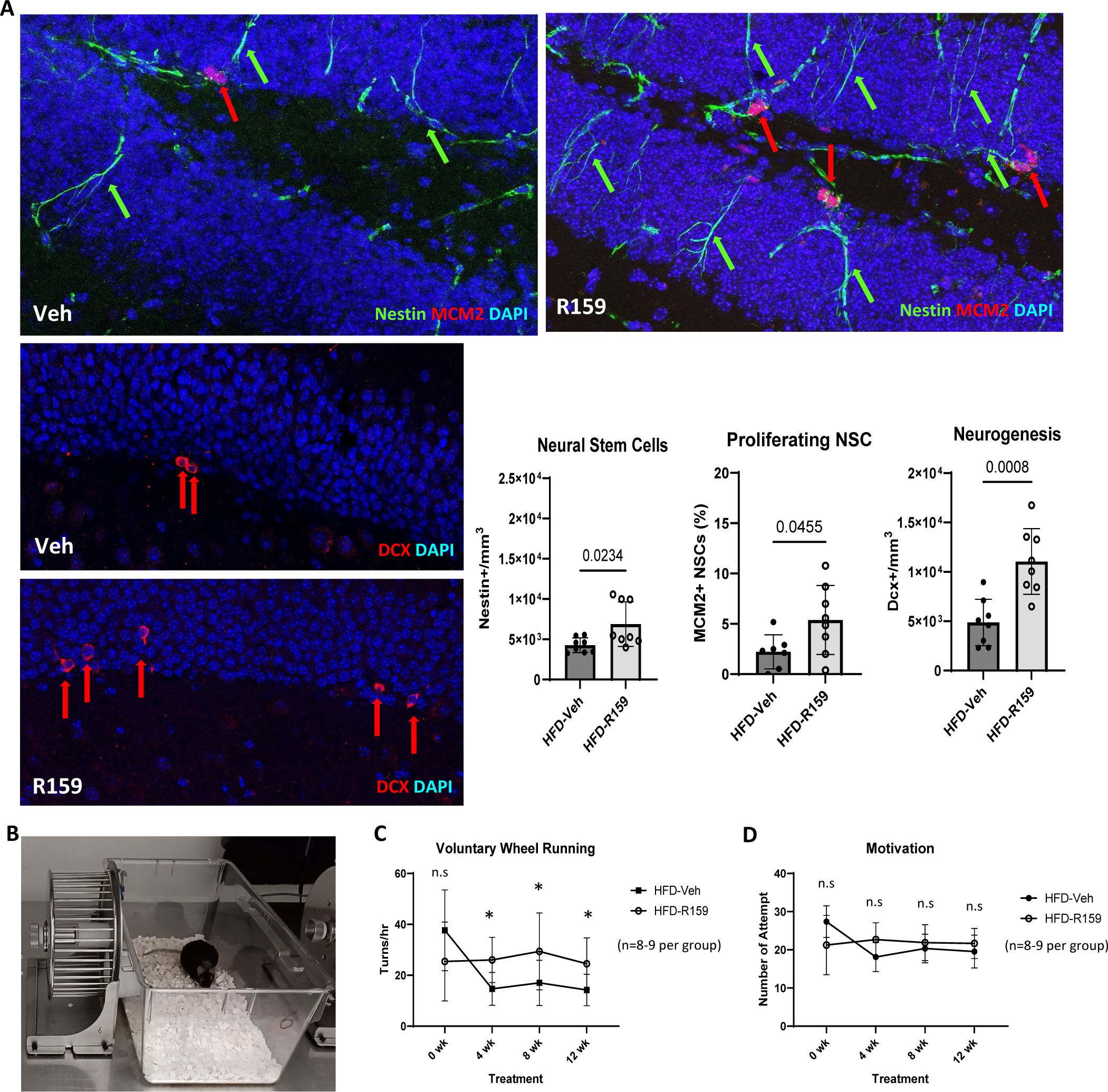
Inhibition of gp130/SFK signaling protects inflammation-induced neurogenesis suppression. (A) The dentate gyrus of the hippocampal region of brains from 16-month-old WT mice on HFD treated with either Veh or R159 were assessed for neurogenesis. Nestin^+^ cells with radial glial branch are quiescent neural stem cells (NSCs); same morphological cells that are Nestin^+^/Mcm2^+^ indicate active NSCs; Doublecortin (DCX^+^) cells mark newly born neurons. Two-tailed Student’s t-test was used for statistical analysis and p-values less than 0.05 were considered significant. (B) Voluntary exercise behavior was assessed through a wheel running assay. (C) Voluntary wheel running activity was measured before and during the drug treatment to assess activity levels. Multiple unpaired t-tests with FDR = 5% were used for the statistical analysis, q-values less than 0.05 were considered significant. (D) Motivation for voluntary exercise behavior was evaluated by counting the number of attempts. Multiple Mann-Whitney tests with FDR = 5% were used for the statistical analysis, q-values less than 0.05 were considered significant.

### R159 treatment increased activity levels in mice on HFD

Observed structural and molecular changes with gp130^Y814^ inactivation demonstrated that the effects of chronic systemic inflammation can be mitigated in multiple tissues in mice on HFD. We investigated whether those changes resulted in a functional improvement. Reduction in activity levels are highly associated with aging and obesity in both animals and humans (38–40). We measured the voluntary activity levels of mice on HFD treated with either R159 or vehicle via a wheel running assay (Figure 5B). R159-treated mice maintained their activity levels despite aging and weight gain, in contrast to Veh-treated mice, which exhibited a significant decline in activity levels (Figure 5C). Interestingly, motivation levels were not different between the two groups, suggesting that R159-treated mice could endure longer or faster runs, indicating less frailty (Figure 5D).

## Discussion

Recent literature highlights gp130 and similar major receptors as complex multimodal sensors, not mere on/off switches, finely tuning diverse signaling pathways (41–43). The findings suggest that selectively modulating a non-canonical gp130 signaling presents a unique therapeutic potential by preserving beneficial canonical signaling pathways, while offering a broader impact than specific receptor blocking by affecting all IL-6 family cytokine signaling. This approach could prove especially beneficial in addressing systemic chronic inflammation associated with aging and obesity, where a delicate balance between inflammation resolution and maintaining essential physiological functions is crucial.

The gp130 receptor primarily activates the JAK/STAT3 signaling pathway which is crucial for regulating immune responses, skeletal development, metabolic homeostasis, and regeneration (44–46). In our recent study, we have identified a novel non-canonical signaling pathway activated by tyrosine (Y) 814 residue within the gp130 receptor that is responsible for the activation of inflammation and fibrosis by SRC kinases (18). We have demonstrated that this signaling can be genetically and pharmacologically inactivated independently of the canonical arm of gp130-JAK/STAT3 pathway and resulted in therapeutically beneficial outcomes in several injury models (18). The receptor modulation strategy, rather than a complete blockade, is fundamentally different from the existing therapeutic approaches. For example, monoclonal antibodies for IL-6, such as tocilizumab, not only block both canonical and non-canonical downstream signaling, but they also have no effect on other IL-6 family members, including OSM, IL-11, and CNTF (47,48). Since all IL-6 members utilize gp130 as the signaling co-receptor, gp130 modulation may not only overcome the negative consequences of complete gp130 blockade that are well-documented (49), but also selectively change the signaling induced by all members of this large family of cytokines. This conceptual innovation was further advanced in the current study using a mouse model of HFD-induced model of chronic inflammation and multimorbidity.

The current study demonstrates that inactivation of a non-canonical gp130 Y814/SFK signaling in both F814 mutant and R159 drug-treated mice exhibited a diminished inflammatory response to HFD in adipose tissue and liver, the major metabolic organs. This led to a significantly lower systemic inflammation, as evidenced by reduced MCP-1 and CRP levels in F814 and R159 treated mice compared to WT or Veh treated counterparts. The upregulation of MCP-1 (*Ccl2*) expression in adipose tissue induced by HFD contributes to monocyte recruitment and macrophage infiltration in adipose tissue and the liver (22,50,51). Our data show that the disruption of IL-6/gp130/SFK activation attenuates the upregulation of *Ccl2* along with other typical pro-inflammatory markers such as *Tnf* and *Il-1b* in the adipose tissue. RNA-seq of the liver tissue revealed reduced expression of *Ccr2* and *Ccr5* in both F814 mice and R159-treated mice compared to their respective control groups, indicating decreased recruitment of macrophages. This was further validated by reduced CD68 staining in the liver sections, and significantly lower systemic CRP levels reflect an attenuated liver inflammatory response. CRP production is primarily induced in hepatocytes by IL-6; CRP is a sensitive clinical marker for systemic inflammation and is known to have a positive association with increasing age and BMI, reflecting the low-grade chronic inflammation characteristic of aging and obesity (6,52).

Histological examination and RNA-seq of the livers showed that WT and Veh treated mice fed HFD had not only more macrophages but also more severe liver fibrosis compared to F814 and R159 treated mice. Non-alcoholic fatty liver disease (NAFLD), precipitated by obesity from HFD, is characterized by excess fat accumulation, fibrosis, and increased immune cell presence (53). NAFLD may progress to more severe stages like steatohepatitis, cirrhosis, and hepatocellular carcinoma if left untreated (54), underscoring the clinical importance for effective interventions. Our findings align with existing literature that demonstrates the inhibition of SFK can alleviate liver fibrosis caused by chemical injuries and metabolic stress (55,56), suggesting gp130 Y814 as a promising therapeutic target for NAFLD. While exploring these implications for NAFLD falls beyond our current scope, our results encourage further investigation.

Remarkably, F814 mutant and R159-treated mice were protected against the loss of trabecular bone and the degeneration of cartilage typically prompted by chronic inflammation in aging and obesity. Bone remodeling, a dynamic process involving osteoblasts constructing the bone matrix and osteoclasts breaking it down, can be disrupted by chronic inflammation, often leading to bone density loss (27,57). IL-6 family cytokines influence this process by regulating RANKL, which is pivotal for osteoclast differentiation (31). The current study found that inhibition of gp130 Y814 signaling significantly reduced RANKL transcription upon hyper-IL-6 stimulation *in vitro*, as well as in obesity-driven systemic inflammatory condition *in vivo*. Furthermore, the crucial role of SRC kinases in osteoclast function is demonstrated by osteopetrosis in SRC-deficient mice (58). Therefore, inhibiting IL-6/gp130/SFK signaling provides a dual protective mechanism against bone density loss in a chronic inflammatory condition by reducing RANKL transcription and SFK activity. Additionally, SFK activity is reported to be critical to cartilage degeneration by promoting cartilage matrix degradation and synovial inflammation (33,59,60). Previous studies by our group demonstrated genetic or pharmacologic inhibition of gp130 Y814 signaling enhances cartilage regeneration in post-traumatic osteoarthritis models in mice and dogs (18). While we observed attenuated cartilage degeneration in both F814 and R159 treated mice compared to their respective control groups, the effect was more substantial in F814 groups than R159-treated group. This discrepancy is likely attributed to the limitations of systemic administration affecting the drug’s bioavailability within the synovial capsule, resulting in less effective cartilage protection compared to direct intra-articular injection.

Systemic chronic inflammation suppresses neurogenesis in humans and mice (61), and HFD is reported to exacerbate neurodegeneration in the hippocampus by disrupting blood-brain barrier, especially in aged mice (62). Neurogenesis in the hippocampus continues into adulthood and has functional association with learning, memory, and mental health (63). We observed that 12-month-old WT mice on HFD exhibited a significant reduction in neural stem cell (NSC) proliferation, whereas F814 mice retained a comparable proliferation rate with ND controls. Notably, 16-month-old mice on HFD treated with R159 showed substantially better NSC number, NSC proliferation, and neurogenesis compared to the Veh group. While reduced levels of systemic inflammation may attribute to the protective effect of R159 against neurodegeneration induced by HFD, the striking differences in NSC niche preservation and neurogenesis suggest a potentially unknown role of IL-6/gp130-Y814 signaling in brain health, warranting further investigation.

IL-6 is a major cytokine of the senescence-associated secretory phenotype (SASP), a complex mix of factors secreted by senescent cells that can influence inflammation and tissue remodeling by affecting the surrounding cells and tissue environment (64). Cell senescence and SASP not only increase with aging but also with an increasing BMI, which is why obesity is considered a form of accelerated aging (65). SASP-driven systemic chronic inflammation is associated with dementia, depression, atherosclerosis, cancers, diabetes, and mortality (66). These morbidities often coexist in aging and obesity, making it hard to treat the root cause and multimorbidity-driven systemic chronic inflammation perpetuates a pathological feed-forward loop of inflammaging (34). Eliminating senescent cells, senolytics, has shown promising results in animal models and human tissues by reducing systemic chronic inflammation and diminishing signs of SASP-associated conditions and are currently under active investigation in multiple clinical trials (67). However, it is a challenging task to specifically target senescent cells due to complex anti-apoptotic pathway networks, and as of now, no senolytics have received FDA approval (54). Our findings demonstrate that selective disruption of IL-6/gp130-Y814 signaling significantly reduced the levels of systemic chronic inflammation, thereby alleviating the associated pathological conditions. While this approach does not eliminate SASP components such as IL-6, it may help mitigate the impact of SASP by reducing levels of subsequent systemic inflammation and degenerative changes across multiple tissues. Consequently, this could lead to an improved healthspan, exemplified by an enhancement in physical activity levels.

The current study has some limitations. All data presented are based on C57BL/6J male mice, considering reported sex differences in response to HFD. Female mice are reported to be less vulnerable to HFD induced inflammation, metabolic disorder, and degeneration, including glucose sensitivity, bone density loss, and cognitive decline (68–71). The underlying mechanisms for the reduced susceptibility of females to HFD-induced inflammatory conditions are not fully understood; however, estrogen emerges as a strong candidate for this protective effect, given its role in mitigating inflammation and metabolic stress (72,73). Therefore, estrogen was considered as a potential confounding factor in our investigation of the effects of IL-6/gp130-Y814 inhibition on obesity-driven low-grade chronic inflammation. In addition, although some initial ADME profile of R159 was conducted, no detailed pharmacokinetic or pharmacodynamic studies of the drug were performed in vivo. A detailed pharmacological assessment is required to explore the bioavailability of the drug in the joints, central nervous system, and other locations notoriously difficult for small molecule targeting. These studies will be essential for the optimization of the treatment regimen in the late preclinical studies.

Collectively, this study highlights the potential of selectively targeting SFK signaling downstream of IL-6/gp130 as an effective approach to reducing systemic chronic inflammation. While degenerative changes and tissue senescence are unavoidable consequences of obesity and aging, this study indicates that the related systemic immune responses and inflammation-driven multimorbidity can be therapeutically addressed.

## Acknowledgments

We thank Dr. Thomas Lozito and Dr. Benjamin Van Handel for their helpful input on the manuscript. We thank Tautis Skorka in USC Molecular Imaging Center for μCT scanning and Seth Ruffins in USC Optical Imaging Facility for providing bone density analysis method. We thank Dr. Lei Peng for mouse perfusion and brain dissection training. We thank master’s students, Falisha Nguyen and Sunyoung Jung, for assisting with sample processing and experiments. We thank USC Department of Animal Resource for animal care and diagnostic service (WBC differential). Diagrams in figures were created with BioRender.com.

Research reported in this publication was supported by the Department of Defense grant W81XWH-13-1-0465 (DE). Research reported in this publication was also supported by the National Institute of Dental & Craniofacial Research; National Institute of Aging; and National Institute of Arthritis and Musculoskeletal and Skin Diseases of the National Institutes of Health under Awards R01AR071734, R01AG058624 and U24DE026914 (all to DE). The content is solely the responsibility of the authors and does not necessarily represent the official views of the National Institutes of Health.

## Author contributions

DE and YL conceived the project. DE, YL, and MB designed experiments. YL conducted vehicle/drug treatment. YL, JT, SL, ACD, JM, US, JL, and DG conducted mouse tissue collection and process. YL, JNL, ACD, JM, HT, TM, and FB conducted tissue staining, imaging, and quantification. YL and JT conducted flow cytometry and analysis. YL and SL conducted ELISA assay. YL conducted RNA extraction and qPCR. AS conducted bulk RNA sequencing and analysis. NL and DG evaluated OARSI and synovitis scores. YL conducted bone density analysis. YL conducted *in vitro* experiments and staining. YL and HT conducted activity assay. YL and TM conducted cholesterol assay. JL and RS conducted western blot. YL and DE wrote the manuscript.

## Data Availability

All data is deposited in GEO data base and is available under the accession number GSEXXXX.

## Methods and Materials

### Animals and treatments

All experimental procedures were approved by the Institutional Animal Care and Use Committee of the University of Southern California and met or exceeded the requirements of the Public Health Service/National Institutes of Health and the Animal Welfare Act. A mutant gp130 Y814 mouse (F814) on a C57BL/6 background was generated using CRISPR-Cas9, and C57BL/6 mice were used as wildtype (WT) controls. All animals used in this study were male. F814 mutant and WT mice were fed a 60% high-fat diet (HFD) (Envigo, TD.06414) from 2-months-old for 10 months. Normal diet control mice were fed a standard rodent diet that contains 13% fat (PicoLab, 5053). F814 and WT mice on a normal diet or HFD were sacrificed at 12-month-old. For drug treatment study, the mice were put on a HFD for 5 weeks to induce obesity before drug or vehicle treatment began. Mice were 13-month-old when drug or vehicle treatment began and were continuously put on a high-fat diet during the 12-week drug/vehicle treatment period. Drug or vehicle treated HFD mice were sacrificed at 16-month-old. Drug R159 was dissolved in a 1% w/v Captisol solution in 0.9% saline at a concentration of 3 mg/mL and administered by intraperitoneal injection at 10 µg per g body weight. The vehicle group was injected with a 1% w/v Captisol solution in 0.9% saline by intraperitoneal injection. The injection volume was kept at 200 µL at a time.

### Dual-energy X-ray absorptiometry (DEXA) scanning

Euthanized mice were scanned using iNSiGHT DEXA scanner (OsteoSys). Low energy at 60 kV and high energy at 80 kV, 0.80 mA current, 5 seconds of scanning at each energy level. An embedded software was used to compile the images and calculate the mass and area of fat and bone mineral content.

### ELISA for serum cytokine/protein

Mouse blood was collected from the submandibular vein and left to clot for 2 hours at room temperature before centrifuging for 20 minutes at 2000 x g. Collected serum was flash-frozen in liquid nitrogen and kept in −80°C until the assay. ELISA kit was purchased from R&D system and experiment was performed following the manufacturer’s instruction; Mouse CRP (MCRP00), Mouse IL-6 (M6000B), Mouse CCL2/MCP-1 (MJE00B).

### White blood cell (WBC) differential

Mouse blood was collected from the submandibular vein in EDTA coated vials and kept in ice until the assay. WBC differential was performed by IDEXX BioAnalytics.

### RNA extraction, gene expression, and quantitative real-time PCR

Total RNA was extracted using the RNeasy Mini Kit (Qiagen) following the manufacturer’s protocol. For gene expression analysis, libraries were prepared using Universal Plus mRNA-Seq with NuQuant (Tecan) after RNA quality validation (Agilent Bioanalyzer 2100) and sequenced on a Novaseq SP (Illumina). For real-time PCR, cDNA was generated using the Maxima First Strand cDNA Synthesis Kit (Thermo Scientific). Power SYBR Green (Applied Biosystems) was used for RT-PCR amplification and detection was performed using ViiA7 (Life Technologies) or Step One Plus Real-Time PCR system(Applied Biolsystems). The comparative Ct method for relative quantification (2−ΔΔCt) was used to quantitate gene expression, where results were normalized to *Rpl7* (ribosomal protein L7). Primer sequences used were: housekeeping gene *Rpl7*: forward 5’ ACCGCACTGAGATTCGGATG 3’; reverse 5’ GAACCTTACGAACCTTTGGGC 3’, Ccl2: forward 5’ TTAAAAACCTGGATCGGAACCAA 3’; reverse 5’ GCATTAGCTTCAGATTTACGGGT 3’, *Tnf*: forward 5’ GACGTGGAACTGGCAGAAGAG 3’; reverse 5’ TTGGTGGTT TGTGAGTGTGAG 3’, tnfsf11: forward 5’ CAGCATCGCTCTGTTCCTGTA 3’; reverse 5’ CTGCGTTTTCATGGAGTCTCA 3’.

### Picro Sirius Red staining

Mouse livers were dissected and fixed in 4% PFA overnight. Tissues were embedded in paraffin and cut at a thickness of 5 µm. Deparaffinized and rehydrated sections were stained with Picro-Sirius Red Stain Kit (Abcam, ab15068) according to the manufacturer’s instructions. Stained sections were imaged using a Zeiss Axio Imager.A2 Microscope Axiocam 105 color camera at three random locations. Positive staining was quantified using ImageJ and an average of three images per sample was used for analysis.

### Immunohistochemistry (IHC)

Mouse tissues were dissected and fixed in 4% PFA for 24 h at 4 °C. Mouse legs were decalcified with 14% EDTA, pH 7.4, for 14-21 days at 4 °C before paraffin embedding. Tissues were embedded in paraffin and cut at a thickness of 5 µm. Antigen was retrieved by Tris HCL pH 10 (Sigma) at 95 °C. Sections were blocked with 2.5% normal horse serum for 1 h at room temperature and incubated with primary antibody (CD68, Invitrogen, PA5-178996) in 1% BSA at 4 °C overnight. Slides were washed and incubated at room temperature for an hour in secondary antibody-HRP (Vector Laboratories, MP-7401). Antibodies were then visualized by peroxidase substrate kit DAB (Vector Laboratories, SK-4100). Slides were imaged using a Zeiss Axio Imager.A2 Microscope Axiocam 105 color camera with Zen 2 program at three random locations. Positive stain was quantified using ImageJ and an average of three images per sample was used for analysis.

### Micro-computed tomography (μCT) data collection and analysis

Mouse legs were dissected and fixed in 4% PFA for 24 h and scanned with XT H 225 ST (Nikon) in 20 µm resolution, at 80 kV energy, 120 μA current, and 9.6 W power at USC Molecular Imaging Center. Trabecular bone under the growth plate near the proximal tibia was reconstructed and analyzed using Amira (Thermo Fisher Scientific). 1 mm length of tibia bone mineral density was measured by bone volume over total volume (BV/TV).

### Flow Cytometry

Flow cytometry was performed on a BD FACS Aria II cell sorter. For adipose tissue immune cells, populations of interest based on negative DAPI (Fisher Scientific, 5748) expression and surface marker expressions Ly6C, Ly6G, CD11b, and F4/80 (BioLegend, 128005, 127639, 123131, 101216) were used. For hematopoietic stem cells, bone marrow cells were stained using BD Stemflow™ Mouse Hematopoietic Stem and Progenitor Cell Isolation Kit (BD, 560492) following the manufacturer’s instructions. For skin stem cells, populations of interest based on negative DAPI (Fisher Scientific, 5748) expression and surface marker expressions ITGA6 (Invitrogen, 12-0495-82), ITGB1(BioLegend, 102215), CD34 (Invitrogen, 11-0341-82), Sca1 (BioLegend, 122513) were used (74). Flow cytometry data was analyzed using FlowJo software.

### TRAP staining and quantification

Mouse legs were dissected and fixed in 4% PFA for 24 h, then decalcified with 14% EDTA (pH7.4) for 21 days at 4 °C. Tissues were embedded in paraffin and cut at a thickness of 5 µm. Deparaffinized and rehydrated sections were incubated in 50 mL TRAP staining medium (110 mM Sodium Acetate (Sigma, S-2889), 50 mM Tartaric Acid (Sigma, T-6521), 4.5 mM Naphthol AS-BI Phosphate (Sigma, N-2125) in distilled water) for 1 hr at 37°C. After incubation, 1 mL of Pararosaniline Dye in Sodium Nitrite solution is mixed into the medium and incubated for 30 min at 37°C. Sections are counter stained with 0.08% Fast Green for 90 sec. To quantify TRAP staining, images were taken from three different locations per sample. Slides were imaged using a Zeiss Axio Imager.A2 Microscope Axiocam 105 color camera with Zen 2 program at three random locations. Positive stain was quantified using ImageJ and an average of three images per sample was used for analysis.

### OARSI scoring and synovitis scoring

Mouse legs were dissected and fixed in 4% PFA for 24 h, then decalcified with 14% EDTA (pH7.4) for 21 days at 4 °C. Knee joint tissues were embedded in the sagittal plane in paraffin and cut at a thickness of 5 µm. For osteoarthritis grade evaluation, deparaffinized and rehydrated sections were stained in 0.4% Toluidine Blue solution for 10 min at room temperature and counterstained in 0.02% Fast Green for 3 min. Research Society International (OARSI) scoring performed as described previously(75). For synovitis grade evaluation, deparaffinized and rehydrated sections were stained in Mayers Hematoxylin for 1 minute and counterstained in alcoholic-Eosin for 1 minutes. H&E staining and synovitis scoring performed as described previously(76). Observers performing the analysis were blinded from the sample IDs.

### Mesenchymal stem cell (MSC) culture and osteogenesis

F814 and WT bone marrow cells were cultured for 10 days in Mesencult^TM^ Expansion Kit for mouse (Stemcell tech, 05513) following the manufacturer’s instructions. After MSC expansion, cells were plated at 10,000/well in a 24-wells plate. When cells reach 80% confluency, cells were cultured in Complete Osteogenesis Differentiation Medium (StemPro, A1007201) for 14 to 20 days with or without 20 ng/mL hyper IL-6 (R&D, 8954-SR) or OSM (R&D, 495-MO) treatment.

### Alizarin Red S staining and quantification

Cultured osteocytes in 24-well plate were fixed in 10% (v/v) formaldehyde at room temperature for 15 minutes and incubated in 40mM Alizarin Red S at room temperature for 20 minutes with gentle shaking. The wells were washed four times with water and imaged by REVOLVE microscope (ECHO). After imaging, 10% (v/v) acetic acid was added (200 μL for 24-well plate) and incubated at room temperature for 30 minutes with gentle shaking. All contents in the well including the monolayer were transferred to a 1.5-mL microcentrifuge tube with a wide-mouth pipette. The tube was incubated at 85 °C for 10 min to extract the dye. After a centrifugation, the supernatant was read in triplicate at 405 nm using Spectramax iD3 (Molecular Devices).

### Osteoclastogenesis and quantification by TRAP staining

Mouse bone marrow was flushed from femur and tibia and plated in 10% FBS α-MEM media supplemented with 30 ng/mL M-CSF (PeproTech, 315-02) for 3 days and then cultured in 10% FBS α-MEM media supplemented with 100ng/mL RANKL (PeproTech, 315-11) and 30 ng/mL M-CSF for 6 to 7 days. Differentiated cells were fixed in stained by 10% (v/v) formaldehyde at room temperature for 15 Acid Phosphatase, Leukocyte (TRAP) Kit (Sigma-Aldrich, 387A) according to the manufacturer’s instructions.

### Western blot

Cells were lysed in RIPA Lysis and Extraction Buffer (Thermo Scientific) containing Pierce Halt Protease inhibitors (Thermo Scientific) followed by sonication with a 15-second pulse at a power output of 2 using the VirSonic 100 (SP Industries Company). Protein concentrations were determined by BCA protein assay (Thermo Scientific). Proteins were resolved with SDS-PAGE utilizing 4–15% Mini-PROTEAN TGX Precast Gels (Biorad) and transferred to Trans-Blot Turbo Transfer Packs (Biorad) with a 0.2-μm pore-size nitrocellulose membrane (Biorad). Nitrocellulose membranes were blocked in 5% nonfat milk in 0.05% (v/v) Tween 20 (PBST) (Corning). Membranes were then incubated with primary antibodies p-gp130 Y814 (Evseenko lab, AbClonal), SRC (Cell Signaling, 2123), p-SRC (Cell Signaling, 6943), or Histone H3 (Cell Signaling, 9515) overnight at 4 °C. After washing, membranes were incubated with rabbit IgG-HRP (Thermo Scientific, 31460) for 1 hour at room temperature. After washing, membranes were developed with the Clarity Western ECL Blotting Substrate (Biorad). Quantification was performed using ImageJ.

### Voluntary wheel running assay

Mice were individually placed in a cage with access to a wheel (Panlab/Harvard apparatus, LE904) for 24 hours. The wheel movement was tracked continuously and recorded automatically in 30-minute increments by a multicounter (Panlab/Harvard apparatus, LE3806). The total number of wheel turns was used to assess activity levels, and the number of attempts was used to assess motivation levels.

### Brain tissue processing and staining

Mice were deeply anesthetized with isoflurane and then transcardially perfused with 10 unit heparin PBS. The brains were isolated and fixed overnight in 4% PFA, followed by incubation in 30% sucrose solution for a subsequent 48 hours prior to sectioning. Frozen coronal sections in 45-µm thickness through the entire dentate gyrus were performed using a sliding microtome (SM2010R, Leica, Wetzlar, Germany). The sections were then transferred to a cryoprotectant solution (27.3% sucrose, 45.5% glycerol, 27.3% ethylene glycol) and stored at −20°C until processing for immunohistochemistry staining.

Immunohistology was performed with antibodies as previously described (77). Briefly, coronal sections of the hemisphere were stereologically sampled throughout the entire hippocampus (5∼6 sections per dentate gyrus), mounted on SuperFrost Plus slides (Thermo Scientific), dried overnight, rinsed in PBS, incubated in 0.01 mol/L citric buffer (pH6.5) for 40 min at 95∼98°C, and rinsed again in PBS. Subsequently, the sections were incubated 2 overnights at 4°C with primary antibodies against the following antigens: Goat anti-Nestin (R&D, AF2736), mouse anti-MCM2 (BD Laboratories, 610701), rabbit anti-DCX (Cell Signaling Technology, 4604S). Sections were then washed with PBS and incubated in appropriate secondary antibodies conjugated to fluorophores (Jackson Immunoresearch) with DAPI counterstaining (Roche, 10236276001). Stained sections were washed, air dried, and coverslipped with PVA/DABCO.

### Quantification of NSCs and Neurogenesis

Images of the stained sections were acquired as a tiled z-stack across the area and the depth containing the dentate gyrus region using a confocal microscope system (Axio.A1 Observer with LSM700 Scanhead, Carl Zeiss, Germany) at 40X. Morphological and co-labeling analysis was done using ZEN 2012 SP1 (black edition, Carl Zeiss, Germany). Nestin+ only cells with radial glial branch are quiescent neural stem cells, same morphological cells that are Nestin+/Mcm2+ indicate active neural stem cells, and DCX+ only cells repregent neurogenesis. The neural stem cell activation rate percentages were calculated through dividing the number of active neural stem cells by the number of total neural stem cells (both Nestin+/Mcm2- and Nestin+/Mcm2+ cells). The total neural stem cell numbers and neurogenesis data were quantified by cells /mm^3^.

