## Supplemental Figures for "Inactivation of a non-canonical gp130 signaling arm attenuates chronic systemic inflammation and multimorbidity induced by a high-fat diet"

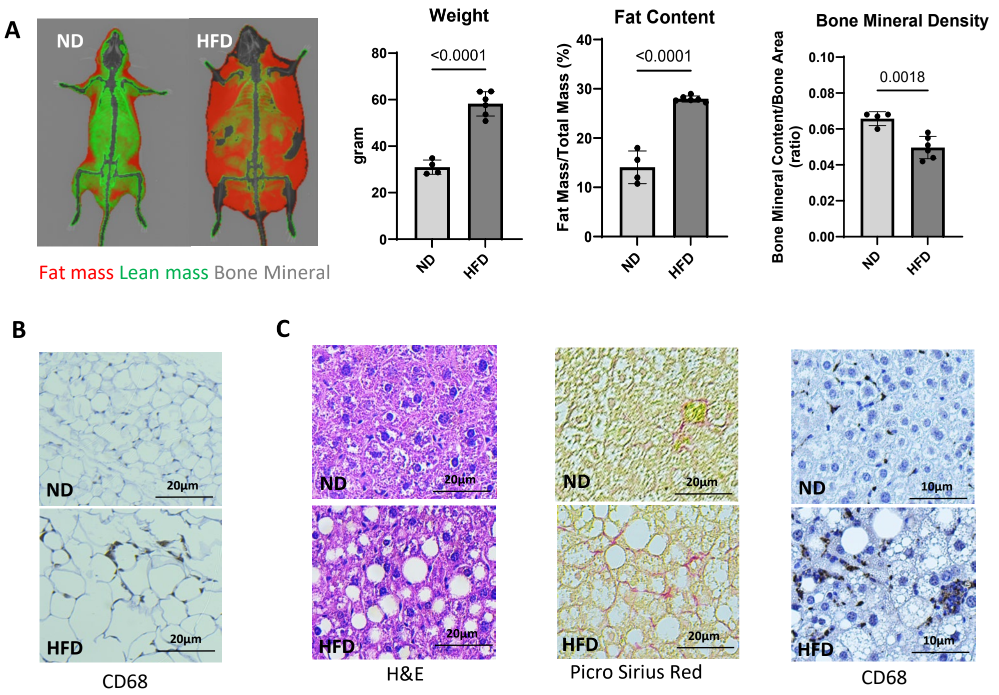


**Figure S1. High-fat diet (HFD) induced changes in wildtype (WT) mice.**
(A) Representative image and body composition analysis of WT mice on a normal diet (ND) or an HFD, generated by Dual Energy X-ray Absorptiometry (DEXA). Fat mass is shown in red, lean mass in green, and bone mineral in grey. Two-tailed Student's t-test was used for statistical analysis and p-values less than 0.05 were considered significant. (B) Representative image of CD68 staining in adipose tissue from WT mice on ND or HFD. Adipose tissue from HFD-fed mice exhibits adipose hyperplasia and necrosis, along with the formation of crown-like structures by macrophages. (C) Representative image of the liver from WT mice on ND or HFD. H&E staining highlights excess fat accumulation indicative of liver steatosis; Picro Sirius Red staining reveals fibrosis; and CD68 staining demonstrates increased macrophage infiltration.

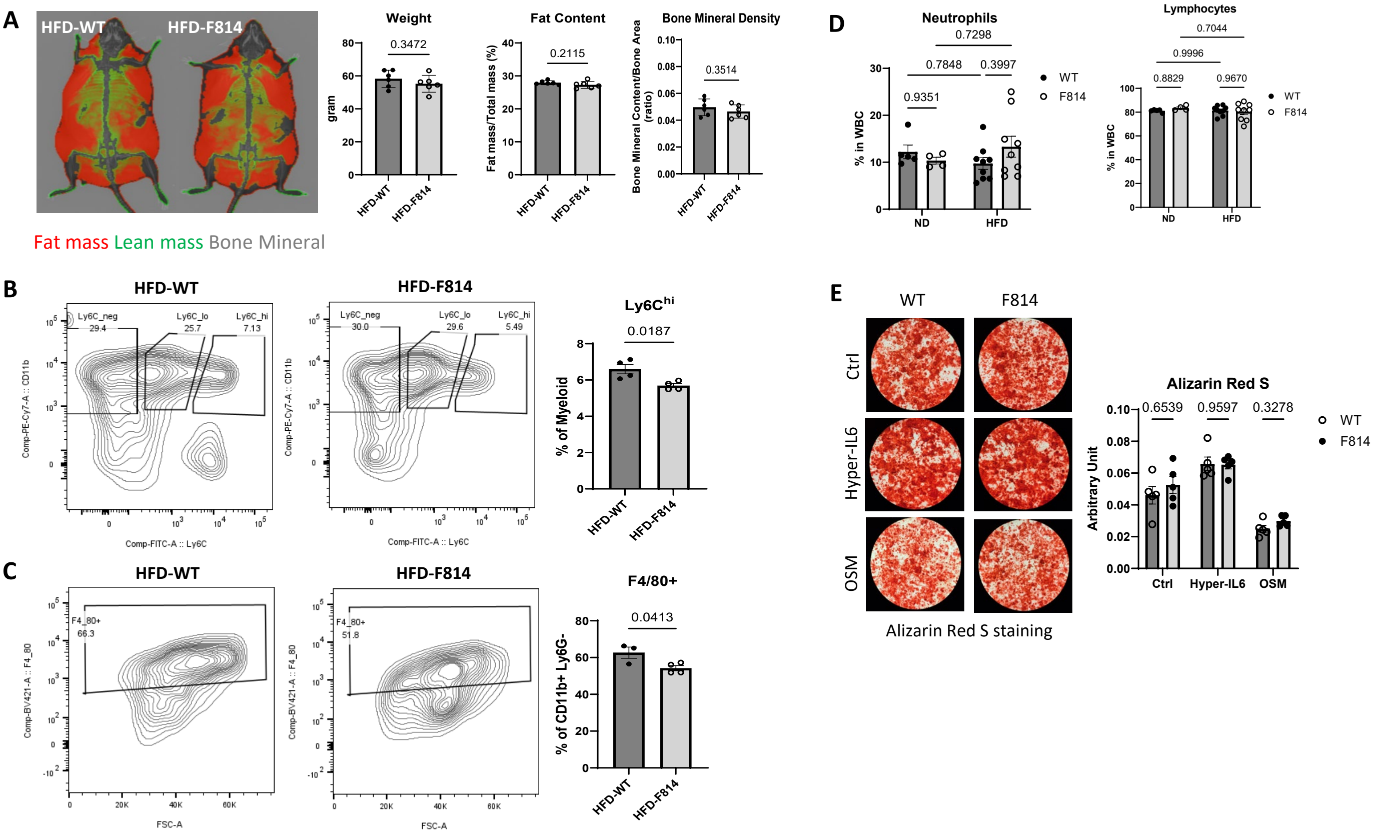


**Figure S2. Impacts of F814 mutation on body composition, immune cells, and bone matrix.**

(A) Representative image and body composition analysis of WT and F814 mutant mice on HFD generated by DEXA. Fat mass is shown in red, lean mass in green, and bone mineral in grey. Two-tailed Student’s t-test was used for statistical analysis and p-values less than 0.05 were considered significant. (B) Flow cytometry analysis of immune cells in adipose tissue from HFD fed WT and F814 mice. CD11b+/Ly6C$^{hi}$ marks pro-inflammatory monocytes. Two-tailed Student’s t-test was used for statistical analysis and p-values less than 0.05 were considered significant. (C) Flow cytometry analysis of immune cells in adipose tissue from HFD fed WT and F814 mice. F4/80+ cells in CD11b+/Ly6G- population marks macrophages. Two-tailed Student’s t-test was used for statistical analysis and p-values less than 0.05 were considered significant. (D) Neutrophil and lymphocyte ratio in peripheral blood analyzed from white blood cell (WBC) differential. 2-way ANOVA with multiple comparisons with Tukey correction was used for statistical analysis and p-values less than 0.05 were considered significant. (E) Alizarin Red S staining of bone matrix produced by osteocytes differentiated from F814 or WT bone marrow derived mesenchymal stem cells and quantification. Multiple unpaired t-test with FDR = 5% was used for statistical analysis and p-values less than 0.05 were considered significant.

**A** Absorbance Distribution Metabolism Excretion (ADME)

| CX ID | TPSA/cLogP/ cLogD/pKa | RLM/HLM %LBF | KS @pH7.4 (µM) | MDCK WT $P_{app}$ A-B/B-A/EER | MDCK MDR1 $P_{app}$ A-B/B-A/EER |
|---|---|---|---|---|---|
| 805 | 55, 3.1, 3.0, 2.1 | 82/89 | 5 | <5/2/1.8 | 6/8/1.4 |
| 159 | 73,0.6,0.5,5.3 | 37/62 | 193 | 102/72/0.7 | 110/75/1 |

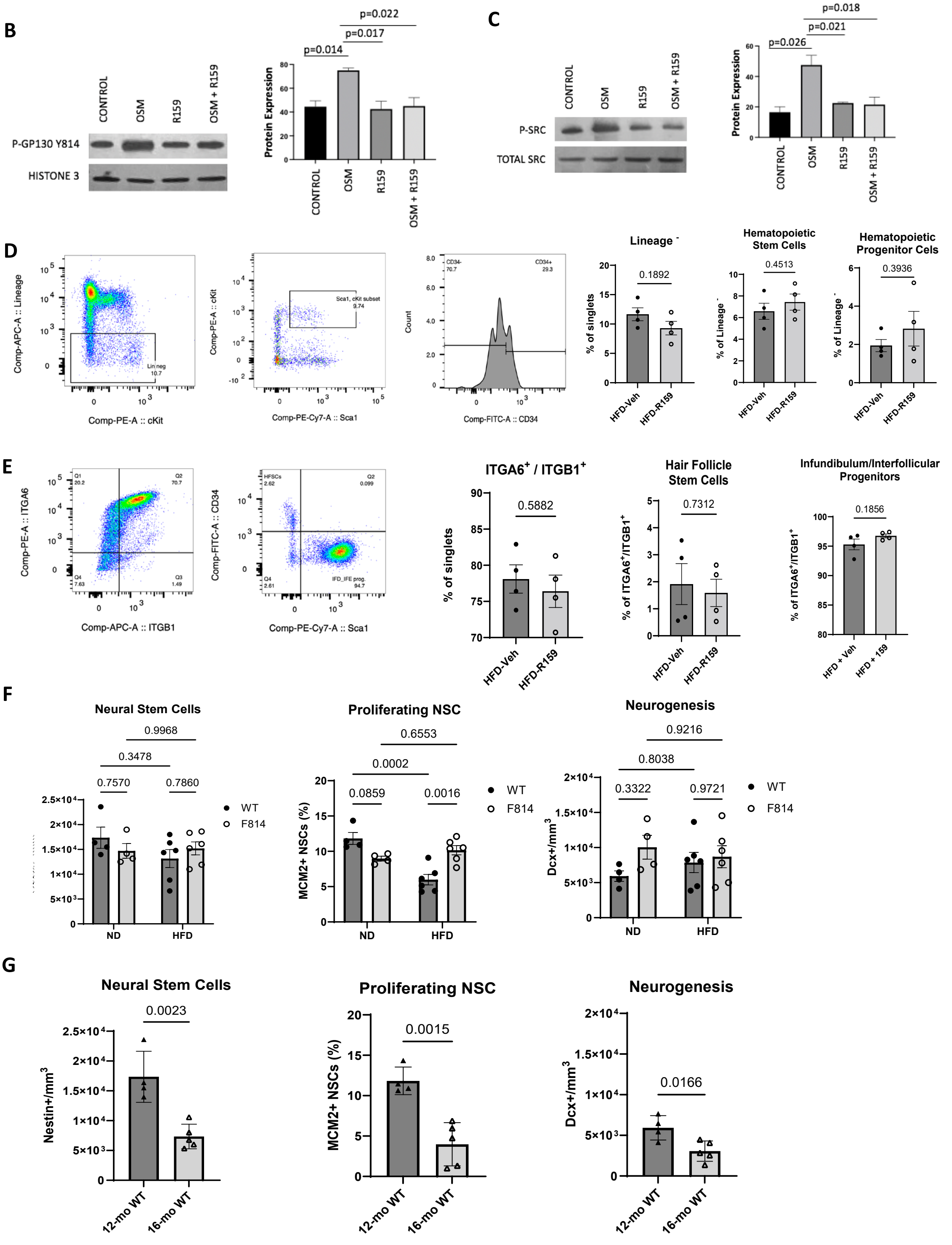

**Figure S3. Gp130 targeting small molecule drug R159 and its effect on aged mice on HFD.**

(A) ADME profile of drugs R805 and R159. TPSA (total polar surface area) range 60-140 Å² is considered good solubility and permeability. cLogP and cLogD show predicted water solubility, cLogP less than 3 and cLogD less than 1 are considered favorable for water solubility. pKa indicates the ionization state of the compound at given pH. RLM/HLM show stability in rat/human liver microsomes, %LBF shows the clearance attributed to liver blood flow. KS shows the solubility constant at a pH 7.4. MDCK WT and MDCK MDR1 cells are used to test drug transport and permeability. A higher Papp value in the A-B direction suggests better absorption potential and a higher Papp value in the B-A direction can indicate strong efflux activity. EER shows an efflux ratio, an EER equal to or close to 1 suggests balanced influx and efflux, indicating that efflux is not significantly hindering the compound's permeability. (B) Western blots for phosphorylated gp130 tyrosin814 in human synovial fibroblast treated with or without OSM (10ng/mL) and R159 (10 µM) for 4 hrs, n = 4. (C) Western blots for phosphorylated SRC in human synovial fibroblast treated with or without OSM (10ng/mL) and R159 (10 µM) for 4 hrs, n = 4. (D) Flow cytometry analysis of bone marrow from 16-month-old C57BL/6 mice on HFD treated with vehicle or R159. The lineage cocktail contains CD3 (a marker for T cells), CD45R(B220) (a marker for B cells), Ly6C and Ly6G (Gr1) (a marker for granulocytes), CD11b (Mac1) (a marker for macrophages), and TER-119 (a marker for red blood cells). Hematopoietic stem cells are marked by Sca-1$^+$/c-Kit$^+$(CD117)/CD34$^{-/dim}$ and hematopoietic progenitor cells are marked by Sca-1$^+$/c-Kit$^+$(CD117)/CD34$^+$. (E) Flow cytometry analysis of skin cells from 16-month-old C57BL/6 mice on HFD treated with vehicle or R159. Hair follicle stem cells are marked by ITGA6$^+$/ITGB1$^+$/CD34$^+$/Sca1$^-$ and infundibulum/interfollicular epidermis keratinocyte progenitors stem cells are marked by ITGA6$^+$/ITGB1$^+$/CD34$^-$/Sca1$^+$. (F) The dentate gyrus of the hippocampal region of brains from 12-month-old F814 and WT mice on normal diet (ND) or a high fat diet (HFD) were assessed for neurogenesis. Nestin$^+$ cells with radial glial branch are quiescent neural stem cells (NSCs); same morphological cells that are Nestin$^+$/Mcm2$^+$ indicate active NSCs; DCX$^+$ cells mark newly born neurons. 2-way ANOVA with multiple comparisons with Tukey correction was used for statistical analysis, p-values less than 0.05 were considered significant. (G) The dentate gyrus of the hippocampal region of brains from 12-month-old WT mice and 16-month-old WT mice on a normal diet was assessed for NSC maintenance and neurogenesis in the same manner as described in (F).

Statistical analyses for (B), (C): 2-way ANOVA with multiple comparisons with Tukey correction, p-values less than 0.05 were considered significant; (D), (E), (F), (G): Two-tailed Student's t-test, p-values less than 0.05 were considered significant.
